# 3D stochastic simulation of chemoattractant-mediated excitability in cells

**DOI:** 10.1101/2021.02.17.431608

**Authors:** Debojyoti Biswas, Peter N. Devreotes, Pablo A. Iglesias

## Abstract

During the last decade, a consensus has emerged that the stochastic triggering of an excitable system drives pseudopod formation and subsequent migration of amoeboid cells. The presence of chemoattractant stimuli alters the threshold for triggering this activity and can bias the direction of migration. Though noise plays an important role in these behaviors, mathematical models have typically ignored its origin and merely introduced it as an external signal into a series of reaction-diffusion equations. Here we consider a more realistic description based on a reaction-diffusion master equation formalism to implement these networks. In this scheme, noise arises naturally from a stochastic description of the various reaction and diffusion terms. Working on a three-dimensional geometry in which separate compartments are divided into a tetrahedral mesh, we implement a modular description of the system, consisting of G-protein coupled receptor signaling (GPCR), a local excitation-global inhibition mechanism (LEGI), and signal transduction excitable network (STEN). Our models implement detailed biochemical descriptions whenever this information is available, such as in the GPCR and G-protein interactions. In contrast, where the biochemical entities are less certain, such as the LEGI mechanism, we consider various possible schemes and highlight the differences between them. Our stimulations show that even when the LEGI mechanism displays perfect adaptation in terms of the mean level of proteins, the variance shows a dose-dependence. This differs between the various models considered, suggesting a possible means for determining experimentally among the various potential networks. Overall, our simulations recreate temporal and spatial patterns observed experimentally in both wild-type and perturbed cells, providing further evidence for the excitable system paradigm. Moreover, because of the overall importance and ubiquity of the modules we consider, including GPCR signaling and adaptation, our results will be of interest beyond the field of directed migration.

**Author summary:** Though the term noise usually carries negative connotations, it can also contribute positively to the characteristic dynamics of a system. In biological systems, where noise arises from the stochastic interactions between molecules, its study is usually confined to genetic regulatory systems in which copy numbers are small and fluctuations large. However, noise can have important roles when the number of signaling molecules is large. The extension of pseudopods and the subsequent motion of amoeboid cells arises from the noise-induced trigger of an excitable system. Chemoattractant signals bias this triggering thereby directing cell motion. To date, this paradigm has not been tested by mathematical models that account accurately for the noise that arises in the corresponding reactions. In this study, we employ a reaction-diffusion master equation approach to investigate the effects of noise. Using a modular approach and a three-dimensional cell model with specific subdomains attributed to the cell membrane and cortex, we explore the spatiotemporal dynamics of the system. Our simulations recreate many experimentally-observed cell behaviors thereby supporting the biased-excitable hypothesis.

## Introduction

How cells sense and interpret external chemoattractant cues and use this information to direct cell movement is one of the most fundamental processes of biology. Unicellular organisms rely on this mechanism to seek nutrients and survive. In multicellular organisms, it is a fundamental process during embryonic development [1, 2] as well as responsible for the proper operation of the mammalian immune system [3, 4]. Perversely, it is through the development of directed cell migration that cancer cells become metastatic [5–7].

Mathematical models have been fundamental in elucidating the mechanisms that cells use to direct the cell migration [8–10]. There is a broad consensus that cells such as the social amoeba *Dictyostelium discoideum* and mammalian neutrophils sense the chemoattractant gradient through a local excitation, global inhibition (LEGI) mechanism based on an incoherent feedforward loop motif that was originally proposed to explain perfect adaptation [11–13]. By incorporating different diffusion properties on the signal components, the mechanism senses static spatial gradients without movement [12, 13].

In response to a spatially uniform stimulus, cells display an initial transient response in which Ras and downstream PI(3,4,5)P_3_ and F-actin activities increase and decrease several times, before eventually returning close to the pre-stimulus basal states, resulting in “near-perfect” adaptation. Signaling motifs responsible for perfect adaptation fall into one of two broad classes: a negative feedback (NFB) loop with a buffering node, or an incoherent feedforward (IFF) loop with a proportioner node [14]. In *Dictyostelium*, experimental evidence favors the presence of IFF-based adaptation [12, 15–17]. The LEGI mechanism, a form of IFF, assumes the existence of a fast local excitor and a slow globally diffusive inhibitor. The productions of the excitor and the inhibitor are independently driven by the receptor occupancy [13]. A local response regulator, which is activated by the excitor and inhibited by the inhibitor, drives downstream signals.

LEGI mechanisms explain gradient sensing but do not account for several aspects of the chemotactic cells, including the ability to move in the absence of external cues. Excitable systems recreate many of the observed properties of randomly migrating cells [18–23] including the stereotypic nature of pseudopods during migration, as well as the spatial pattern of activities exhibited by both signaling and cytoskeletal elements in cells [24, 25]. Recently, LEGI mechanisms have been coupled to an excitable system to explain how a cell’s ability to migrate randomly can be steered in the direction of the external gradient [17, 22]. When combined with a memory-like ability to polarize the chemotactic machinery, these models account for nearly all the observed behavior of chemotaxing cells [22, 26]. Most mathematical models of chemoattractant signaling have adopted a variant of the FitzHugh-Nagumo (FHN) model of neuronal excitability [27, 28]. These models, however, suffer from limitations because of the phenomenological aspect of the model. For example, using reaction terms in polynomial form as in the FHN model does not permit direct biological interpretability. Moreover, system states, which would typically represent concentrations, are not constrained to be nonnegative [19–22, 29].

Though the LEGI mechanism successfully explains the temporal and spatial responses to chemoattractant in *Dictyostelium* [15–17, 30], little attention has been paid to the effect of noise on the LEGI mechanism. Moreover, in the excitable paradigm, the ability to generate random protrusions, as seen in unstimulated migrating cells, relies on stochastic fluctuations triggering the excitable system. A proper account of these fluctuations is vital, however, as it their relative size that determines whether the external signal directs movement properly. In practice, noise arises as an intrinsic feature of the stochastic nature of the biochemical reactions and depends on the state of the dynamical system [31, 32]. To our knowledge, however, all existing computational models use partial differential equations and generate these fluctuations by injecting noise as an additional input into these differential equations.

To overcome the aforementioned limitations, here we present a new model of the biological signaling mechanism that regulates motility. Specifically, to account for the noise accurately we eschew the partial-differential equation approach and instead incorporate the reaction-diffusion processes into the URDME (Unstructured Reaction Diffusion Master Equation) software [33]. This methodology does not make a priori assumptions on the size of the stochastic perturbations; instead, the fluctuations are inherent in the underlying chemical master equation. Hence, the noise is controlled by number of the molecules of the reacting species. Moreover, we consider a realistic geometry consisting of a three-dimensional cell with membrane and cortex elements. Finally, we use detailed biochemical models of the receptor dynamics, LEGI and excitable modules and verify the sufficiency of these three interconnected networks in explaining the spatiotemporal dynamics of chemotactic signaling.

## Methods

### Overall system architecture

For the present study we have divided the signaling pathways that drive the observed excitable Ras dynamics in *Dictyostelium* cells into three signaling subsystems (Fig. 1A): 1) The G-protein coupled receptor (GPCR) subsystem is responsible for sensing chemoattractants. 2) The receptor occupancy information from GPCR is passed down to the Local Excitation and Global Inhibition (LEGI) mechanism which is responsible for adaptation in the presence of a global stimulus as well as interpreting directional cues when cells are in a chemoattractant gradient. 3) The Signal Transduction Excitable Network (STEN), accounts for the features of the excitable behavior, including all-or-nothing responses, refractory periods and wave propagation, and provides a characteristic response both in the presence or absence of chemoattractant signals. We have excluded downstream networks that directly influence the actin dynamics. The output of our system could readily be coupled to the cytoskeletal signaling network but that would likely necessitate numerous additional entities, including elements with a large number of molecules (e.g., ~10^8^ molecules of actin [34]) leading to a presently unmanageable computational burden for these stochastic simulations.

**Fig 1.**
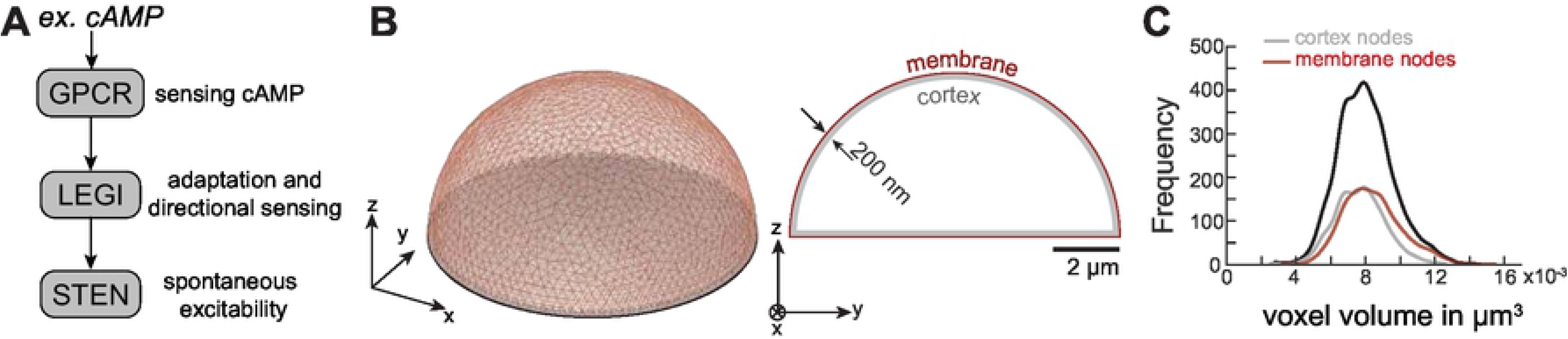
Model schematic and simulation domain. (A) Reaction scheme adopted in the present study involves a receptor module describing GPCR (G-Protein Coupled Receptor) dynamics, a LEGI (Local Excitation and Global Inhibition) module that provides adaptation and directional sensing, and a STEN (Signal Transduction Excitable Network) that describes the excitable behavior of the cell. (B) Isometric (left) and cross-sectional side view (right) of the hemispherical simulation domain of radius 5 *μ*m and thickness of 200 nm. The outer surface is the membrane and the interior of the shell is the cortex.(C) Frequency distribution of all the nodal volumes (black), membrane nodes (red) and cortex nodes (gray).

### Geometry

We used a stationary hemispherical structure of radius 5 *μ*m to capture the general shape of adherent amoeboid cells devoid of any cytoskeletal components (as if treated with Latrunculin) (Fig. 1B). Because most of the species known to be involved in generating the spatial patterns are either membrane-bound, or in the cytoskeletal cortex, we focus on these areas of the cell. We do not incorporate nodes for the cytosol but consider this a sink from some molecules can move in and out of. We assumed that the cortex is 200 nm thick and discretized a shell of this thickness into nodes resulting from an unstructured tetrahedral mesh. The allowable minimum and maximum distances between the nodes were set at 100 and 300 nm, respectively. Overall, this gave rise to 5,500 nodes and 16,469 tetrahedral elements with volume distribution 26.8 ± 4.6 × 10^−4^ *μ*m (mean±std. dev.). Because a coarser mesh affects the smoothness of diffusion, a finer mesh would be preferred, but this increases the simulation cost considerably and would become a major constraining factor as the number of biochemical elements in the model grew (S1 FigA–D). For this reason, we compromised by choosing a medium size mesh so that, in an element of average size, a single molecule corresponds to a concentration of approximately 60 nM. The nodal volumes obtained from the dual of the tetrahedral mesh has a leptokurtic distribution with parameters: 80.1 ± 16.3 × 10^−4^ *μ*m (mean±std. dev.). The nodes on the surface are denoted as the membrane nodes whereas the others are assigned as cortex nodes (Fig. 1C). The volume distributions of cortex and membrane nodes were similar, with means of 8.4 × 10^−3^*μ*m and 7.7 × 10^−3^ *μ*m, respectively. The mean subtracted distributions failed to reject the null hypothesis using a Mann-Whitney-Wilcoxon test, indicating that the two size distributions are not significantly different. The distributions of membranes nodes at the basal and apical surfaces were also similar, with mean volumes of 7.3 × 10^−3^ *μ*m and 7.9 × 10^−3^ *μ*m, respectively.

## Results

### GPCR signaling module

The initiation of chemotaxis in *Dictyostelium* involves binding of chemoattractant molecules (ligand) to surface-bound G-protein coupled receptor (GPCR) molecules. In the present study, we focused on cAR1 as the major receptor for cAMP. The heterogeneous 3’,5’-cyclic adenosine monophosphate (cAMP) binding in *Dictyostelium* has been modeled using three states indicating different levels of affinities [35]. For simplicity, we ignored the sequential binding dynamics and followed the classification of GPCRs on the basis of affinity only. The three states of unoccupied GPCRs have high (H), low (L) and slow (S) binding affinity with respect to extracellular cAMP [35]. The corresponding occupied receptor states are labeled H:C, L:C and S:C, respectively. Interaction of cAR1 with cAMP also results in the desensitization of receptors due to phosphorylation [36, 37]. Despite any notable loss in the binding sites, a 3–5-fold decrease in the binding affinity of the low affinity class (L) has been reported [36]. This motivated us to consider additional phosphorylated receptor states: ^P^H, ^P^L, and respective occupied states: ^P^H:C, ^P^L:C. Our complete receptor model consists of 10 reacting species and 26 reactions in total (Fig. 2A). The detailed reactions and parameter values in mesoscopic form are listed in Table 1.

**Fig 2.**
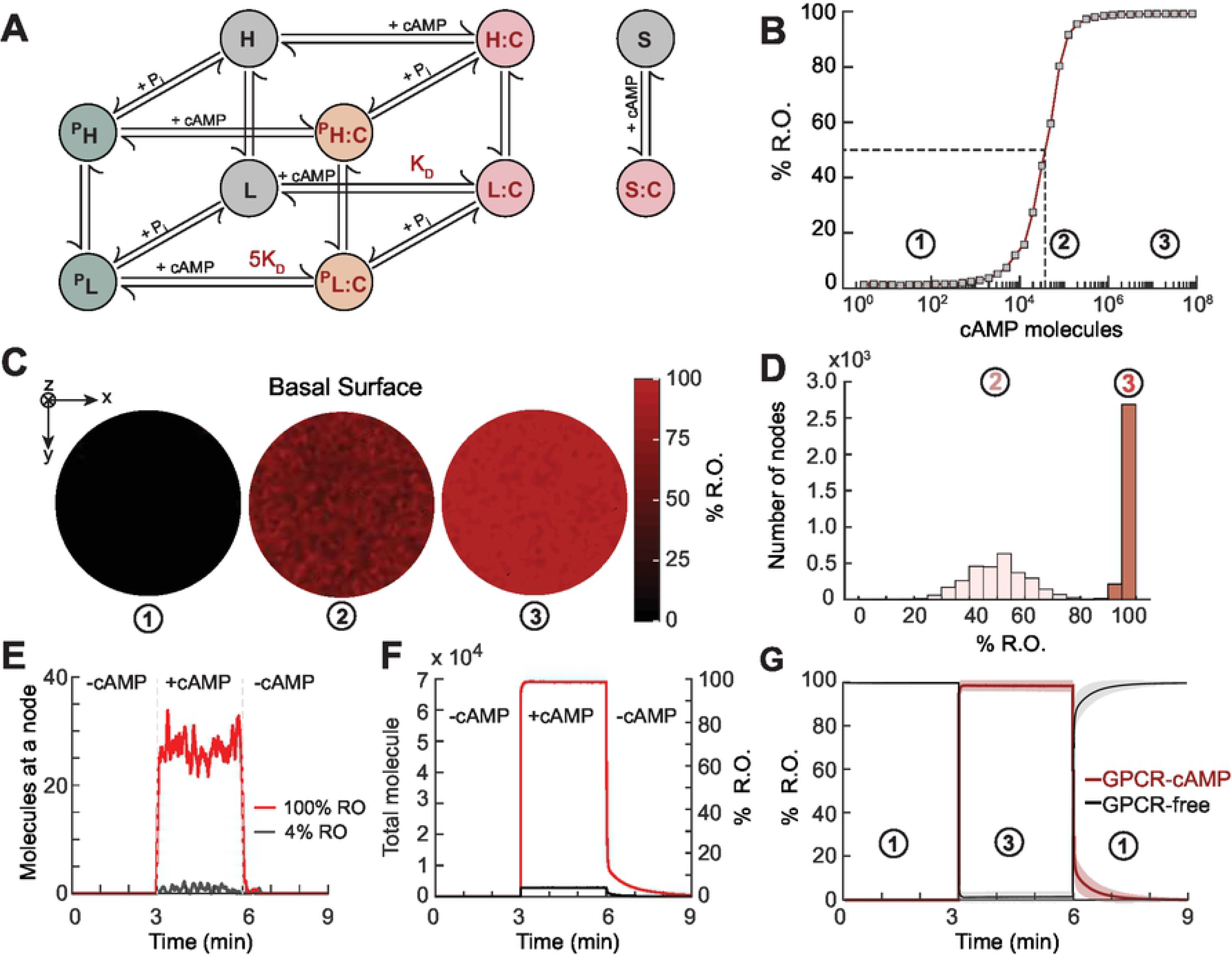
GPCR signaling. (A) Detailed schematic of the different states of the G-protein coupled receptor (GPCR) and cAMP binding. Unoccupied receptors exist in high (H) and low (L) affinities, and a third slow (S) binding state. Occupied receptors are denoted H:C, L:C and S:C. Phosphorylated states are denoted by a superscript P. (B) Dose response curve. The circled numbers denote different concentration levels of cAMP corresponding to (1) low (4%), (2) mid (50%) and (3) high (100%) levels of R.O.(C) Steady-state R.O. in response to different concentrations of cAMP.(D) Distribution of nodes based on R.O. for mid (light red) and saturating (red) cAMP doses.(E,F) Temporal profile of number of total occupied receptors (H:C+L:C+S:C+^P^H:C+^P^L:C) at a single random node (E) and in the cell (F) for low (black) and saturating (red) doses of cAMP.(G) Temporal profile of total free (black) and occupied (red) receptors in a cell in response to application and removal of the high dose of cAMP. The shaded regions denote the respective standard deviations (*n* = 10 independent simulations in which the parameter values are allowed to vary according to the distributions of 1).

**Table 1.**
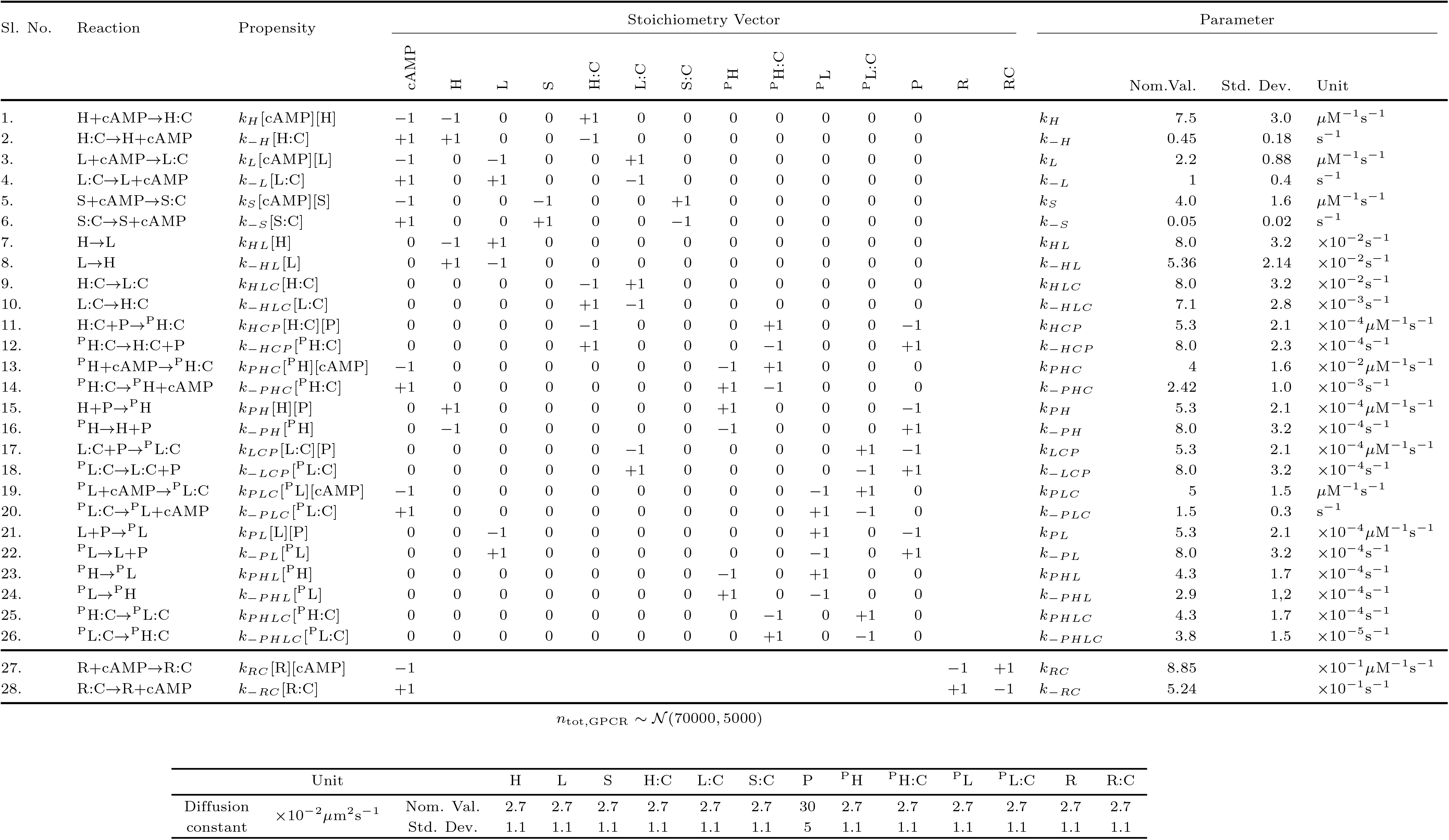
Parameters for G-Protein-Coupled Receptor (GPCR) Module with propensity and stoichiometry vectors. Parameter values for reactions (1–10) are from [35]; for reactions (11–26) the values were estimated to match the experimental observations from [36, 37]; for reactions (27–28) the values were obtained by minimizing the difference between the responses of full- and reduced-order system. The diffusion constants for GPCRs are from [38]

To study the system response to different levels of cAMP, we mapped the relationship between the dose of the cAMP (in terms of the total number of molecules of cAMP experienced by the cell) and receptor occupancy (R.O.) (Fig. 2B). We considered three different cAMP doses corresponding to low (4%), mid (50%, EC50) and high (100%, saturating dose) levels of R.O.. The respective profiles at the basal surface show little variation at the two cAMP concentration extremes, but considerably more heterogeneity (skewness = 0.04) at the mid-point (Fig. 2C,D). In this case, the individual nodes displayed an approximately normal distribution, but some nodes (*n* = 4) had as few as 10% or as high as 85% R.O. (*n* = 2). With higher cAMP concentrations, the distribution became more skewed (skewness = −1.77), but there was a small number of nodes (*n* = 18) with as little as 90% R.O. (Fig. 2D).

In the presence of cAMP, the free GPCR states get converted to the respective occupied states, among which the occupied low affinity receptors (L:C) form the greater population (S2 FigA,B). The phosphorylated states showed slower dynamics (*t*_1/2_ = 198 s) compared to the unphosphorylated states (*t*_1/2_ ~ 10 s). As might be expected from a mostly Poisson process (average Fano factor = 0.95, S2 FigC) the absolute value of the temporal fluctuations at individual nodes increased with increasing cAMP concentrations but the fluctuations in the fraction of occupied receptors decreased (Fig. 2E). These trends held when considering fluctuations in the sum of all nodes, though the relative noise became smaller (Fig. 2F). Furthermore, we looked into the heterogeneity in terms of the mean occupied receptor states among nodes for a range of doses of cAMP. We observed that the relative internodal noise in the system decreases with the increasing dose of cAMP and eventually approaches a Poisson distribution (average Fano factor = 0.85) for the saturating dose of cAMP from the initial sub-Poissonian distribution (S2 FigD,E). In a population of cells, the number of receptors and other signaling molecules differs from cell to cell. To examine this heterogeneity, we simulated receptor occupancy in models in which the cells had varying number of receptors as well as allowing the parameters to vary (Fig. 2G, S1 File). We observed differences (std. dev. ~2.5% at the saturating dose, and ~10% during initial decay phase following removal of stimulus) in the receptor occupancy level among cells when subjected to the same cAMP dose.

Lastly, because of the computational burden of dealing with the large number of states and reactions, we considered the possibility that GPCR binding could be represented by a reduced-order model that could capture the essential spatiotemporal dynamics as well as most of the noise characteristics of the full model. To this end, we used a model consisting of one free (R) and a single occupied (R:C) state (S2 FigF) and varied the parameters of the reduced model so as to minimize the error between the temporal response of the full and reduced systems. The responses matched closely with only small differences during the initial decay phase following the removal of the stimulus (S2 FigG,H). Importantly, the noise characteristics were also similar over a wide range of cAMP doses; for example, at a saturating dose of cAMP, the standard deviation of the full order was 4.58 molecules compared to 4.59 for the reduced-order model. (S2 FigC,E). This finding suggests that there is little error in using the more computationally efficient reduced-order model in lieu of the detailed one.

### LEGI module

Though the LEGI mechanism successfully explains the temporal and spatial responses to chemoattractant in *Dictyostelium* [15–17, 30], little attention has been paid to the effect of noise on the LEGI mechanism. Moreover, there are a number of possible ways of implementing LEGI, several of which we considered. In *Dictyostelium*, chemoattractant signaling depends on the presence of G-proteins. G-proteins form a heterotrimer, consisting of *α* (we focused on G_*α*2_, which mediates cAMP signaling downstream of the cAR1 receptor [39, 40], *β* and *γ* subunits. Whereas these subunits are together in an unoccupied receptor, the *α*2 and *βγ* subunits dissociate upon stimulation at which time the latter are free to signal to down-stream elements. This dissociation is persistent [40]. The G*_βγ_* works upstream of Ras and is crucial in chemotaxis [41, 42]. The Ras response, as well as downstream signals, display properties of excitable systems, including a refractory period [24], an all-or-nothing threshold [43], and wave propagation [44–47]. At the same time, near perfect adaptation is observed in the activated Ras response [16]. This suggests that adaptation happens upstream of Ras and, as the response of G*_βγ_* is persistent during cAMP stimulation, we treated this as the excitation process in the LEGI module (Fig. 3A). The parameters for the G*_βγ_* dynamics were chosen (Table 2) to match experimentally measured half-times of dissociation (on application of a saturating stimulus) and reassociation (on removal of stimulus) of the G_*α*2_ and G_*βγ*_ subunits [17].

**Table 2.**
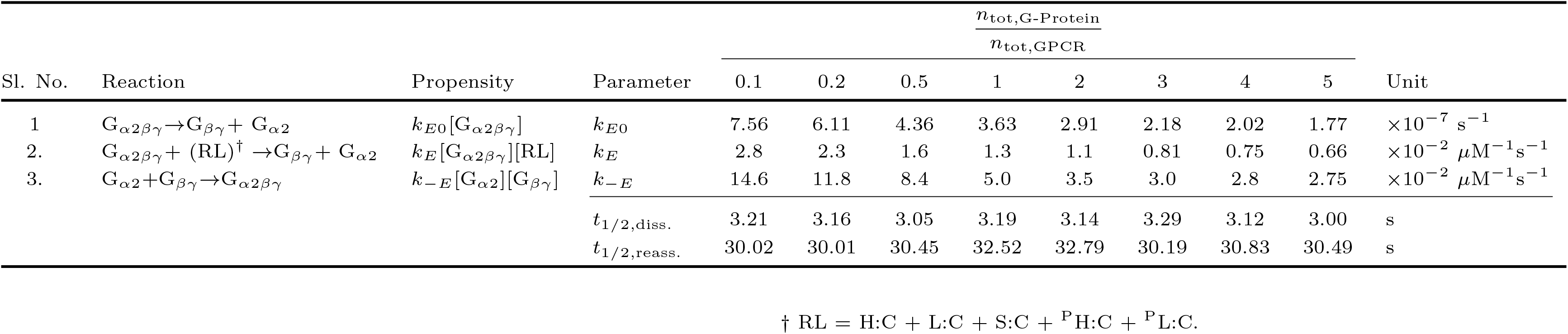
Parameters for G-Protein Dynamics. The parameter values are estimated to match the half times of dissociation (*t*_1/2,diss._) and reassociation (*t*_1/2,reass._) from [17]

**Fig 3.**
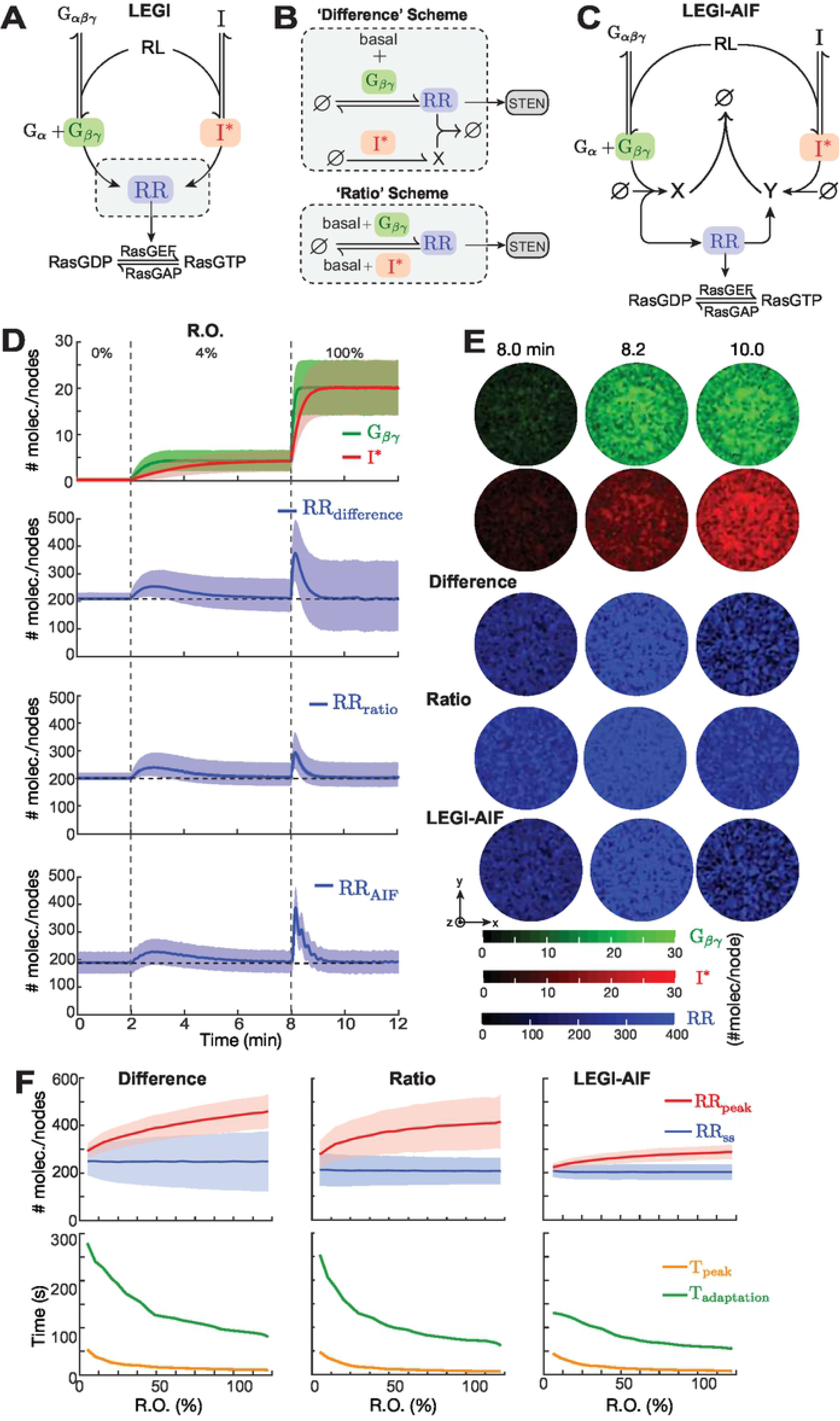
Response of the LEGI mechanism to global stimulation. (A) LEGI scheme involves a local activator (G_*βγ*_) and global inhibitor (I^⋆^). Their interaction creates a response regulator (RR) which positively affects the conversion of RasGDP to RasGTP. (B) The regulation of RR can be realized either through a Difference scheme (top) or through a Ratio scheme (bottom). Both involve basal and G_*βγ*_-dependent production of RR. Whereas inactivation is mediated by I^⋆^ through an intermediate X in the difference scheme, in the ratio scheme it depends on both basal and I^⋆^-dependent terms. (C) LEGI with Antithetic Integral Feedback (LEGI-AIF). In this scheme, G_*βγ*_ and I^⋆^ create intermediates (X and Y, respectively) that annihilate each other. The RR is created by Y and catalyzes the production of X. (D) Temporal dynamics of components of the different LEGI schemes. The top panel shows nodal average profiles of G_*βγ*_ (green) and I^⋆^ (red) in response to a staircase profile of cAMP stimulus: 0–2 min: 0% R.O.; 2–8 min: 4% R.O.; 8–12 min: 100% R.O. The bottom panels show the corresponding RR profile for the different schemes. The shaded regions denote standard deviations among all nodes from a single simulation. (E) Basal surface profile of G_*βγ*_ (green), I^⋆^ (red) and RR (blue) from different schemes at the time points indicated. (F) Effect of concentrations of cAMP (in terms of % R.O.) on peak amplitude (red), steady-state amplitude (blue), peak time (orange) and adaptation time (green) of the nodal average profile of RR for different schemes. The solid lines and the shaded regions show the respective mean and standard deviations (*n* = 10 independent simulations as in Fig. 2G).

We simulated the system and looked at the dissociated subunits and observed a highly nonlinear relationship between the response curves of receptor occupancy and the dissociated G-protein (S3 FigA). The fraction of dissociated G-proteins decreased as the total number of G-proteins increased (S3 FigA); however, as a function of the maximal response, it was independent of the total number of G-proteins reaching 50% dissociation at ~23% R.O. (S3 FigB), indicating the existence of “spare receptors” in the system [48]. Additionally, we observed that fluctuations (mean coefficient of variation) in the G-protein response decreased with higher total number of G-proteins (S3 FigC). We chose the total G-proteins to be three times the total number of receptors as for the higher values there was not difference in the relative fluctuation level.

Unlike the excitation process, inhibition is G-protein independent [17] in *Dictyostelium*. To account for this, we assumed an inhibitor existing in both active (I^⋆^) and inactive (I) forms. The diffusion constants for the inhibitor states were assumed high (a number typical of cytosolic entities [47]) to satisfy LEGI requirements that the global inhibitor have higher diffusivity than the local G_*α*2*βγ*_ and G_*βγ*_. To account for a possible multistep translocation to and from the membrane, we made the inhibitor dynamics slow compared to those of G_*βγ*_.

The interaction of the excitor G_*βγ*_ and the activated inhibitor jointly regulate a response regulator, RR, whose action can be modeled using either a difference or a ratio scheme, depending on how the activation and inhibition processes regulate it (Fig. 3B, S3 FigB–D). In the difference scheme, basal and G_*βγ*_-dependent activation together with I^⋆^-mediated deactivation of RR through an intermediate lead to a steady-state RR concentration that is an affine (linear plus a constant) function of the difference between G_*βγ*_ and I^⋆^ (Suppl. S3 FigC, S1 File). An alternative realization, the ratio scheme, involves G_*βγ*_-dependent activation and I^⋆^-dependent inactivation of RR along with independent basal activation/inactivation and leads to a steady-state of RR that is proportional to the ratio of affine functions of the G_*βγ*_ to I^⋆^) (S3 FigD, S1 File). As there are a number of potential biochemical elements that could serve as the response regulator, we considered both schemes. We implemented these schemes explicitly, using the corresponding reactions, as well as implicitly, using a quasi-steady-state approximation of the reactions. By ignoring the transient dynamics, the implicit formulations served to reduce the computational burden of explicitly simulating these systems (S1 File).

One of the possible limitations of the adaptive networks described above is the sensitivity of the adaptation to noise, resulting in variances that do not adapt, but depend on the stimulus level [49]. As a third alternative, we adopted the Antithetic Integral Feedback (AIF) model [50] by making some of the terms local and global, and then compared its performance with the two schemes described above (Fig. 3B). Here, G_*βγ*_ and I^⋆^) create intermediates that annihilate each other. One intermediate activates RR which, in turn, catalyzes the production of the other. The detailed reactions and parameter values of all the three LEGI schemes are listed in Table 3.

**Table 3.**
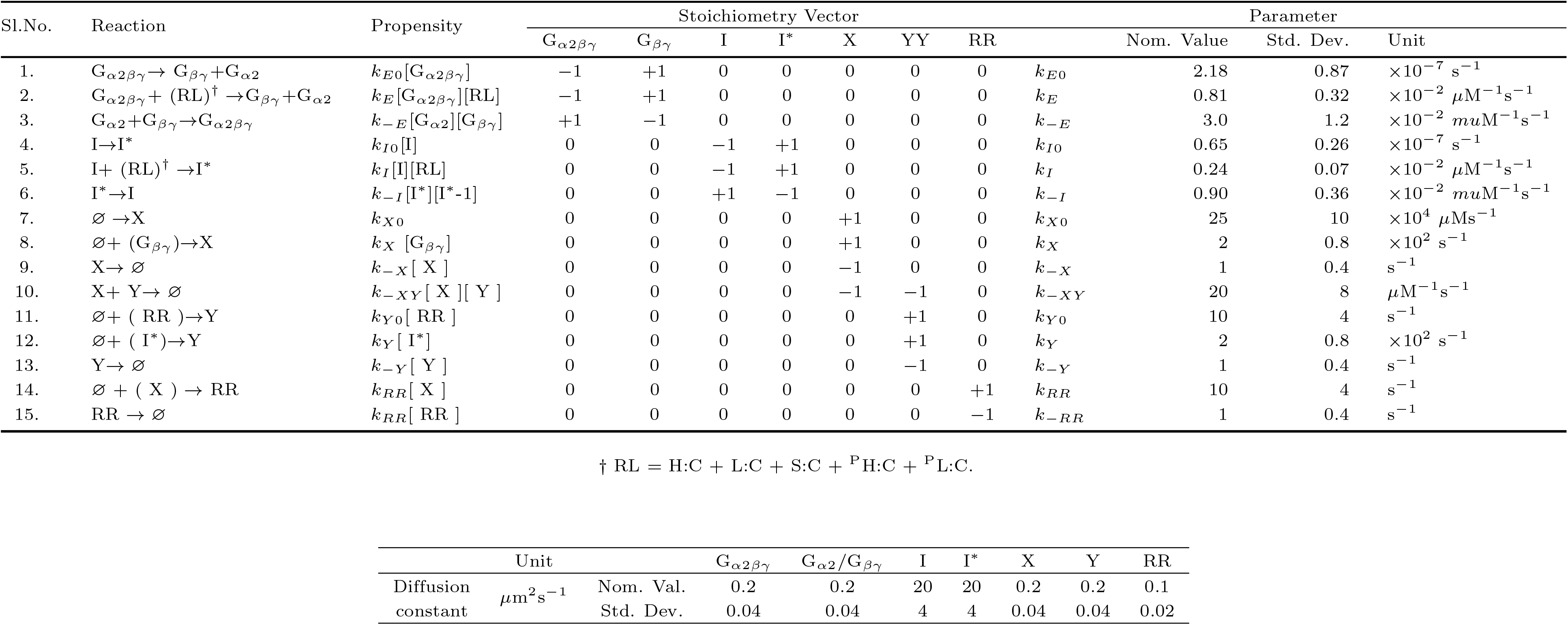
Parameters for LEGI and LEGI-AIF module.

### Simulating the combined GPCR and LEGI modules

To analyze the adaptation characteristics of proposed LEGI networks, we applied patially uniform increasing concentrations of cAMP starting from no stimulus, (0% R.O.; 0-2 min) rising to a low dose (4% R.O.; 2-8 min) and finally reaching a saturating dose (100% R.O., 8-12 min). These simulations were done in models that allowed for varying internal concentrations and parameter values so as to capture cellular heterogeneity. As expected, both G_*βγ*_ and I^⋆^ rose with the latter showing a slower response (Fig. 3D,E; Fig S4A,B). In all variants considered, the initial responses depended on the stimulus level; i.e., the peak amplitude (red) increased and the peak time (orange) decreased with %R.O. (Fig. 3F). In all three networks, the mean nodal response regulator activity returned to prestimulus levels (blue in Fig. 3F). However, the steady-state variance showed a stimulus-level dependency that increased for the difference scheme, decreased slightly for the ratio scheme, and was smallest and fairly constant for the AIF scheme (blue in Fig. 3F). The adaptation (settling) times in all three schemes decreased monotonically as a function of %R.O., in agreement with published experimental data [16], with the AIF scheme showing the least sensitivity (green in Fig. 3F). Unlike the difference and ratio schemes, the temporal profile of the LEGI-AIF showed some oscillatory behavior during the adaptation at high levels of R.O.. The variance in the peak amplitude of the initial response for the ratio scheme was higher than for the difference scheme. The lower coefficient of variation in the peak response profile of the difference scheme provides more certainty of response to change of stimulus than the ratio scheme. Upon removal of the stimulus, the difference scheme returned to the basal level more quickly than to the ratio scheme (S4 FigC,D). For the rest of study, we used the implicit difference scheme of LEGI to avoid unnecessary repetition.

We next simulated the response to a gradient generated by releasing chemoattractant from a micropipette. To this end, we imposed a stationary 3D Gaussian profile centered around an edge point at the basal plane and considered both the apical and basal (Fig. 4A) and perimeter responses (Fig. 4B; S1 File). Following imposition of the gradient, free G_*βγ*_ rose quickly at the front where %R.O. was highest; at the rear the rise was slower (Fig. 4C). In contrast, the inhibitor rose more slowly and was fairly uniform around the perimeter of the cell, owing to the relatively high diffusion (Fig. 4C,D). The resultant response regulator rose sharply at the front and dropped below the basal level at the rear (Fig. 4C,D). Note that the steady-state level of the response regulator at the front was lower than the peak, as the level of the inhibitor increased, but still displayed levels above basal. To examine the role of diffusion in these patterns, we varied the inhibitor diffusion over a wide range (S5 Fig). Increasing the ratio of inhibitor-to-G_*βγ*_ diffusion resulted in greater differences in the response regulator between front and back. The mean difference between RR molecules between front and back doubled for a 10-fold difference in diffusion coefficients.

**Fig 4.**
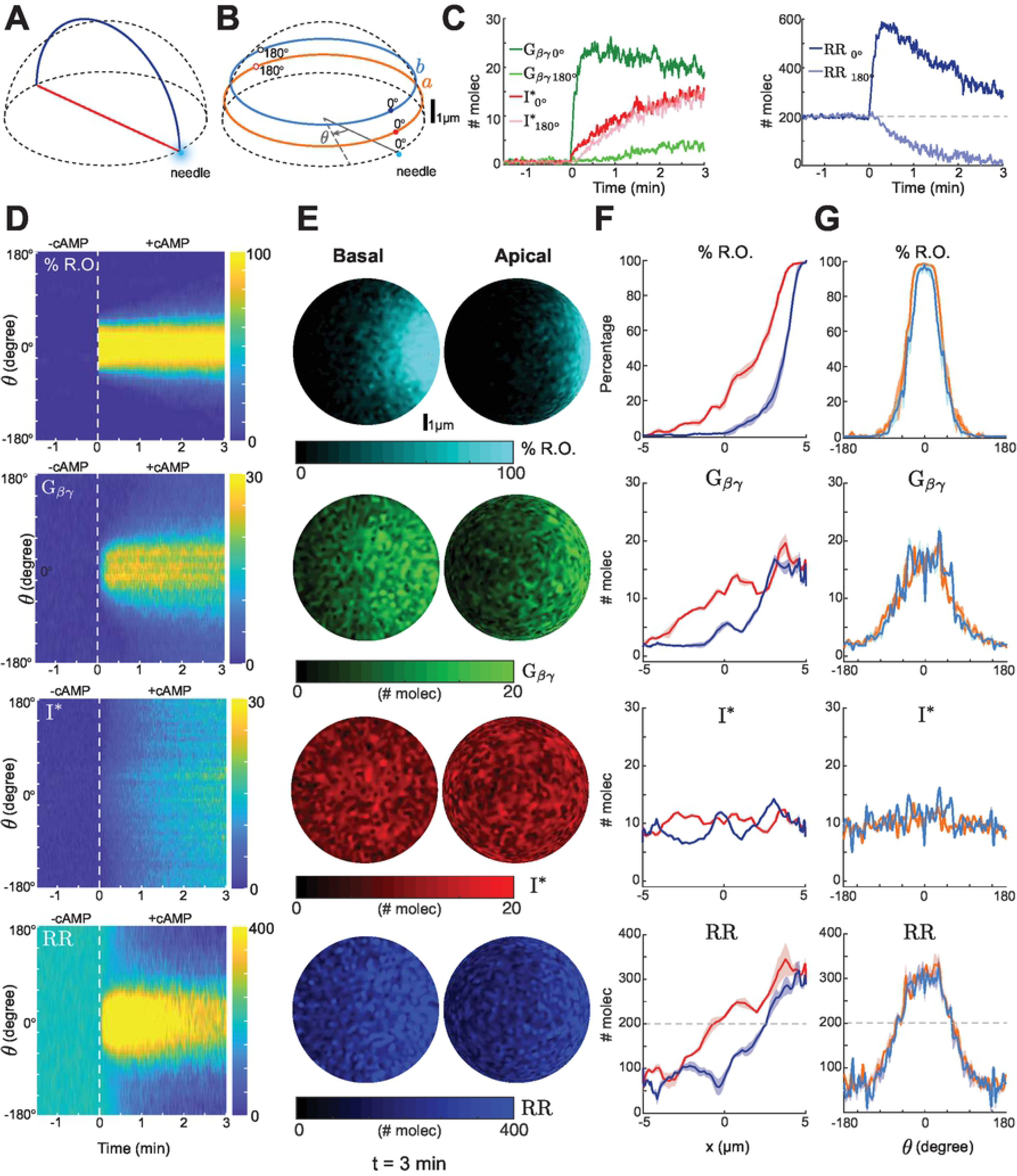
Steep gradient sensing by LEGI difference scheme. (A) Schematic showing the diameter at the bottom surface (red) and a semicircular arc on the curved surface (blue) connecting front and back of the cell with respect to the needle position (light blue dot). (B) Circular cross sections of the hemispherical domain at *z* = 1 (a) and *z* = 2 μm (b). The front (closest to needle) and back of the cell is marked as 0 and 180°, respectively. (C) Temporal profiles of G_*βγ*_ (green), I^⋆^ (red) and RR (blue) at cell front (0°, darker shade) and back (180°, lighter shade). The solid lines and the shaded regions show the respective mean and standard deviations (*n* = 10 independent simulations as in Fig. 2G). (D–G) Spatial response of the system for receptor occupancy (R.O.), G_*βγ*_, I^⋆^ and response regulator (RR). The kymographs (D) are based on the maximal projection of the hemispherical domain (nodes between a and b) of panel B. The white dashed line indicates the time instant when cAMP gradient was applied. Panel E shows the spatial profiles at the basal and apical surfaces at *t* = 3 min. Panel G shows the spatial profiles along the lines marked in pane A. Lines denote mean and the shaded regions standard deviations (*n* = 5 independent simulations as in Fig. 2G).

At steady state, the linear profiles of local entities such as R.O. and G_*βγ*_ displayed gradients that were more diffused on the top surface than on the basal surface (Fig. 4E–G). In contrast, the inhibitor profile was relatively flat along the length of the cell, though random variations were apparent. The resultant response regulator profile showed a gradient and was higher/lower at the front/rear relative to the basal level. We repeated the simulations for a shallower gradient and observed similar behavior, with a smaller degree of localization and slower response in RR (S6 Fig).

### Signal Transduction Excitable Network (STEN) module

*Dictyostelium* cells have two excitable systems that work in tandem: a fast cytoskeleton-based network, and a slower signaling network that drives the cytoskeletal system [24, 25, 44]. As we are modeling cells lacking an intact cytoskeleton, we focused on the latter. Our proposed STEN consists of five species: RasGDP, RasGTP, activated and inactivated protein kinase B substrates (PKB*s and PKBs, respectively), and membrane phosphatidylinositol bisphosphate (PIP_2_), which represents contributions from both PI(3,4)P_2_ and PI(4,5)P_2_ (Fig. 5A).

**Fig 5.**
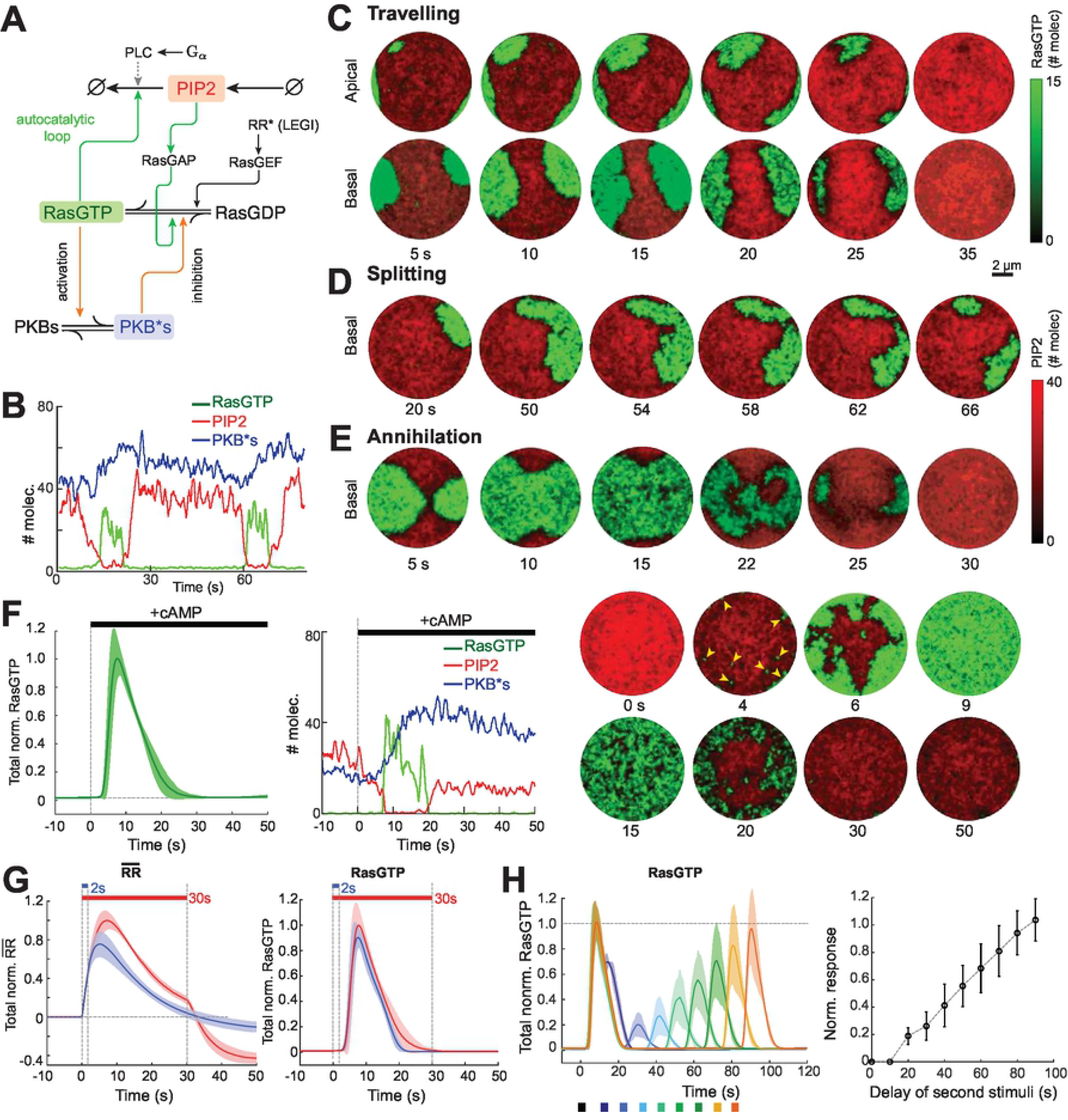
Response of STEN to global stimulus. (A) Schematic of STEN showing the entities: RasGTP, RasGDP, PIP2, PKBs, PKB*s and their interactions. Green arrows denote positive feedback whereas, orange arrows complete the negative feedback on RasGTP. (B) Temporal profiles of RasGTP (green), PIP_2_ (red) and PKB*s (blue) at a random node that has fired spontaneously. (C–E) Spatiotemporal profiles of RasGTP (green) and PIP_2_ (red) of the membrane showing wave-traveling (C), splitting (D) and annihilation (E). (F) Response to a spatially uniform dose of cAMP. Shown are the global RasGTP response (left), various component at a single random node (center) and the spatiotemporal profile at the basal surface of the cell (right). The yellow arrowheads denote wave initiation sites. Colors are as in panels B–E. The shaded region in the left panel is the standard deviation as in Fig. 4D (*n* = 10). (G) Basal subtracted normalized RR (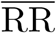, left) and RasGTP (right) responses to short (2 s, blue) and long (30 s, red) stimuli. The solid lines and the shaded regions are as in panel F (*n* = 10). (H) RasGTP response to the two short pulses (2 s) of spatially uniform stimuli with variable delays. Left: temporal mean nodal profile of RasGTP (*n* = 10). Right: plot of normalized peak of the second response to the first versus the delay between the stimuli.

In our scheme, RasGTP acts as the activator of the excitable system and serves as a front marker of the cell. Recent experiments have demonstrated that lowering PI(4,5)P_2_ results in increased Ras activity [43]. Similarly, lowering PI(3,4)P_2_ increased Ras activity through the regulation of RasGAP2 and RapGAP3 [51]. Thus, we incorporated the PIP_2_-mediated hydrolysis of RasGTP to RasGDP into the model. This closes a positive feedback loop that is formed through mutual inhibition between RasGTP and PIP_2_.

A slower, negative feedback loop is achieved through the RasGTP-mediated activation of PKBs. This activation is achieved partly by having PH-domain containing PKBA translocate to the membrane to bind to PI(3,4,5)P_3_, as well as TorC2-mediated phosphorylation of PKBR1 [52]. This negative feedback loop is closed as activated PKBs (PKB*s) negatively regulate RasGTP. There are two proposed mechanisms of negative feedback: 1) controlling the localization of the Sca1/RasGEF/PP2A complex on the plasma membrane through phosphorylation of Sca1; 2) phosphorylation and activation of PI5K which increases PIP_2_ [53, 54]. As we do not specifically model PI(3,4,5)P_3_, we implemented this loop with PKBs being activated directly by RasGTP, with a slower time scale to account for the omitted intermediate steps (e.g., PI3K activation, PI(3,4,5)P_3_ formation and PKB translocation and subsequent phosphorylation) and that some reactions involve cytosolic species, compared to the mutually inhibitory positive feedback loop.

Finally, we coupled this excitable network to the LEGI in two ways. First, through an RR-dependent term that converts RasGDP to RasGTP, consistent with the possibility that the RR is a RasGEF or activates a RasGEF. Second, we also included a term to account for the activation of Phospholipase C (PLC) by G_*α*2_ which leads to PI(4,5)P_2_ hydrolysis [55]. Table 4 lists the detailed reaction terms and parameter values.

**Table 4.**
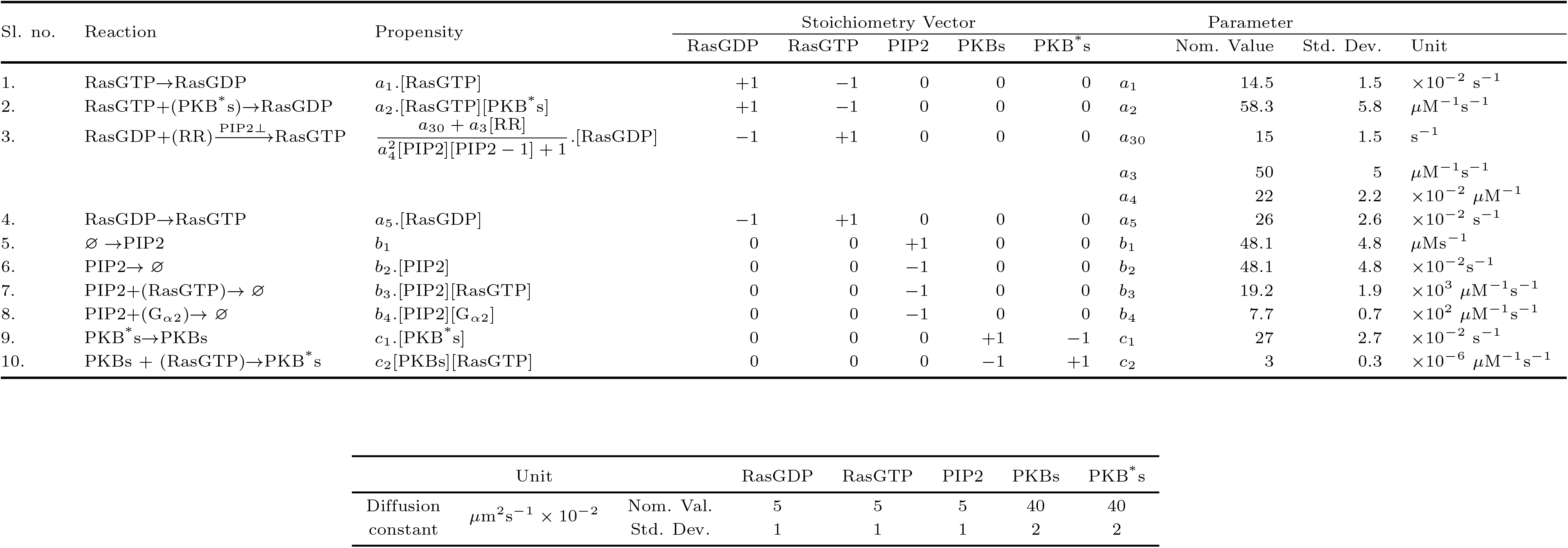
Parameters for Signal Transduction and Excitable Module. Stoichiometry and propensity functions.

### Combining the GPCR, LEGI, and STEN modules

We first simulated our model in the absence of any stimulus. We observed characteristic excitable behavior (Fig. 5B,C). When viewing a single node, RasGTP and PIP_2_ showed mutually exclusive behavior, in which the number of molecules at any one time of one species dominated the other. For example, when PIP_2_ dominated, there was approximated 40 molecules of PIP_2_ compared to approximately one molecule of RasGTP (Fig. 5B, *t* = 30 s). The transition to a state in which RasGTP dominated (~25 molecules of RasGTP vs. ~0-2 molecules of PIP_2_) was rapid. In contrast, the number of PKB*s molecules varied considerably less, ranging from ~40 to ~60 with slow increases happening after the node transition to a high RasGTP state (Fig. 5B, *t* = 60 s). When viewed across the surface of the cells, we observed wave activity (Fig. 5C,S7 FigA,B,S1 Video). The cell was typically in a back state in which high PIP_2_ levels dominated. However, when stochastic perturbations led to a spot of high RasGTP activity, a wave of activation swept across the cell, moving between the basal and apical surfaces. This eventually extinguished and the cell returned to its basal back state.

Interestingly, after the RasGTP wave went through a region, the subsequent PIP_2_ recovery was actually higher than before the wave, indicating an overshoot of the basal level (c.f. the PIP_2_ intensity between the first and last panels in Fig. 5C, S7 FigA (bottom)). The elevated PIP_2_ regions played an important role in the steering of waves. Whenever a traveling wave encountered a region of supra-basal PIP_2_ on its path, it moved around this region which often resulted in the splitting of the wave (Fig. 5D, S2 Video). Generally, wave splits were rare because of the small size of the cell relative to the wave. This is consistent with experimental findings which have prompted experimentalists studying waves to consider giant fused cells [44, 46]. Consistent with properties of excitable waves, when two wave fronts met, they annihilated (Fig. 5E, S3 Video). The annihilation time for wave collisions depended on the time scale and relative diffusivity of RasGTP and PKB*s.

### Response to stimulation

We simulated the response of our model to a spatially uniform stimulation using saturating dose of cAMP (100% R.O., Fig. 5F, S4 Video). After an initial delay of 2–3 seconds, we observed multiple wave initiations (arrowheads in Fig. 5F) and wave spreading. The waves eventually spanned the whole cell at which time (~8–9 s) total Ras activity peaked. The activity died down as the RR level returned to the pre-stimulus condition. The experimentally observed response of cells is quite similar following short or long pulses of chemoattractant stimulation, consistent with the notion that the underlying signaling network is excitable [7, 24]. To test this in our model, we repeatedly applied short (2 s) and long (30 s) global stimuli to models with varying parameter values and compared the respective RR and RasGTP responses (Fig. 5G). The response profiles were nearly indistinguishable, with both peaking about 7–8 s after the application of the stimulus, followed by a return to the pre-stimulus level. They only differed in the final phase in which the response to the short stimulus reached its steady-state faster (~20 vs. ~30 s). Interestingly, the RasGTP response was only noticeable after ~2 s at which time the short stimulus had already been removed. This is consistent with excitable systems which reach a point of no return following stimulation. When the stimulus remained present beyond the time taken for the excitable system to return to its basal state (> 60 s), we observed a smaller and more patchy second wave of activity (S7 FigC–F). This second peak of the response is due to the partially adapted state of the LEGI module which is due to the adaptation time being longer than the duration of a pulse from the excitable system; it has been seen experimentally both with signaling [56] and cytoskeletal biosensors [57].

Cells displays refractory periods following stimulation during which further excitation fails to trigger a response, or diminished in intensity [7, 24]. We applied a series of double pulses, each of duration 2 s with variable delays (10–90 s) and compared the peak of the second response to the first response (Fig. 5H, left panel). For short delays (< 10 s) there was no distinct second response. However, as we increased the time delay, we observed an increase in the peak amplitude of the second response. After a sufficiently long delay (80 s), the second peak almost matched the first peak with a 50% recovery occurring at ~50 s. (Fig. 5H, right panel).

When stimulated by a gradient, cells display persistent high levels of activity at the side of the cell facing the chemoattractant source. We simulated these experiments by introducing a gradient of receptor occupancy across the cell. RasGTP showed a localized persistent patch of high activity facing the side with highest receptor occupancy (Fig. 6A, S5 Video). We also simulated two experiments in which the spatial gradient was combined with a temporal stimulus. In the first, we introduced a gradient and waited until the cell displayed a spatial response; we then removed it for a variable time, before finally reintroducing it [58]. The crescent disappeared following removal of the stimulus but did not return to its full strength until the delay was ~60 s (Fig. 6B, S6 Video), matching our previous observations in the double pulse experiment. In the second simulation, we applied a large global stimulus following the establishment of a crescent in response to a gradient, and then removed all cAMP. In this case, the global stimulus elicited a response everywhere (Fig. 6C, S7 Video). These simulations show the complex interactions of spatial and temporal components of coupled LEGI and STEN systems.

**Fig 6.**
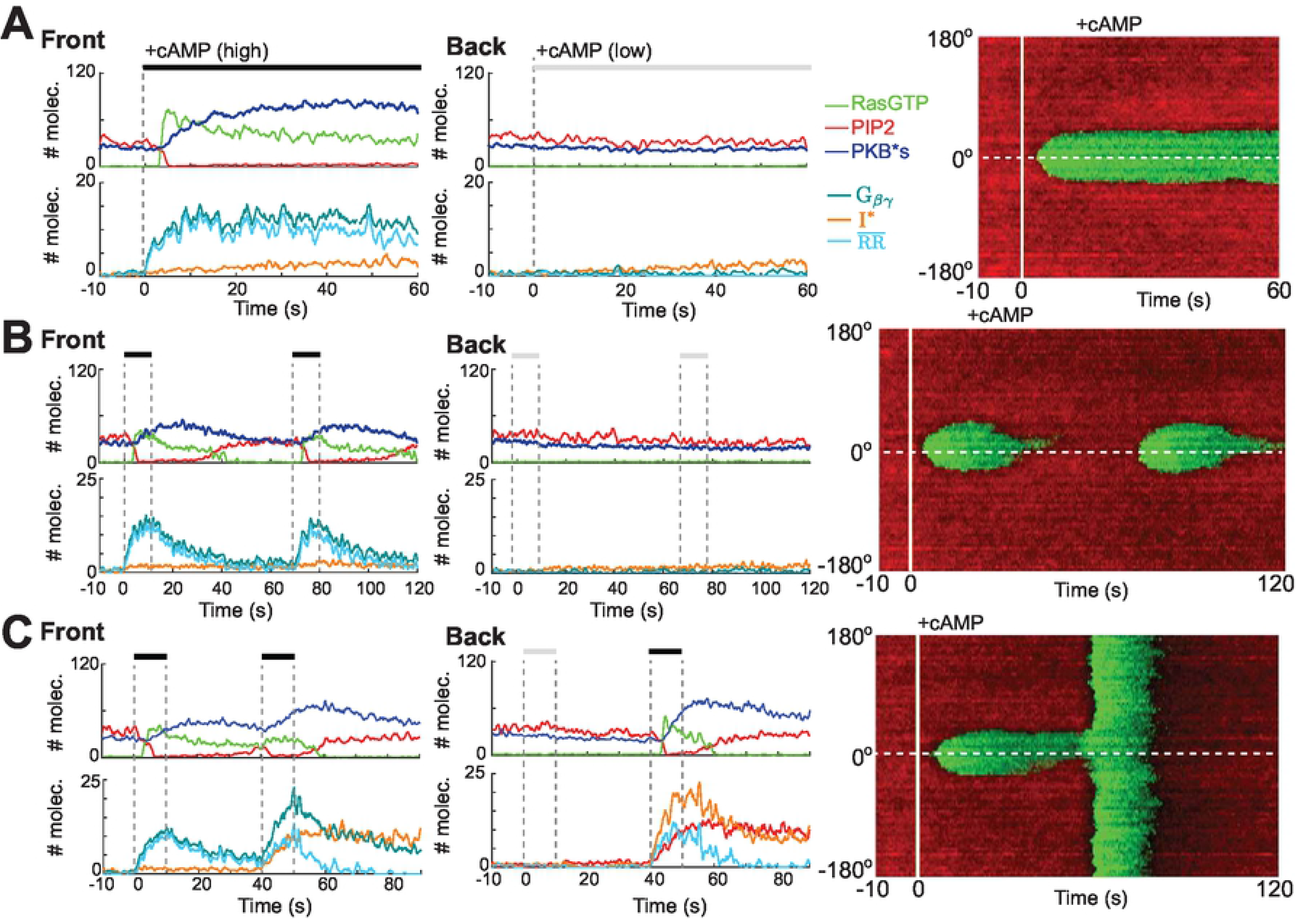
Gradient sensing by STEN. (A–C) Response to temporal and gradient stimuli. Shown are the temporal profiles of RasGTP (green), PIP_2_ (red), PKBs*(blue), G_*α*2_ (teal), G_*βγ*_ (orange) and basal subtracted normalized RR (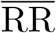, cyan) at single nodes at the front and back of the cell. The kymographs (right) show RasGTP (green) and PIP_2_ (red). Solid white line denotes when the gradient was applied, and the dashed line shows the needle position (front). Whereas Panel A shows the response to a single gradient, B and C show the response to two gradient stimuli with a delay of 60 s (B) and to a gradient stimulation followed by a global one (C).

### Effect of lowering threshold

Recent experiments in which the activity and motility of the cell were altered suggest that these came about through changes in the threshold of the excitable signaling system. Our model allows to test some of these perturbations. We first considered lowering PIP_2_ levels by adding an extra degradation term, so as to recreate experiments in which the phosphatase Inp54p is brought synthetically to the cell membrane [7, 44]. In this case we saw elevated levels of activity and the cell commenced periodic whole-cell increases in activity similar to what have been observed experimentally (Fig. 7A). Similar, though less acute increases in activity were seen in simulations in which PKB*s levels in the cell were lowered through a reduction in the RasGTP-mediated activation rate (Fig. 7B). In this case, bursts of wave initiations were observed, but the resulting waves did not cover the whole cell surface.

**Fig 7.**
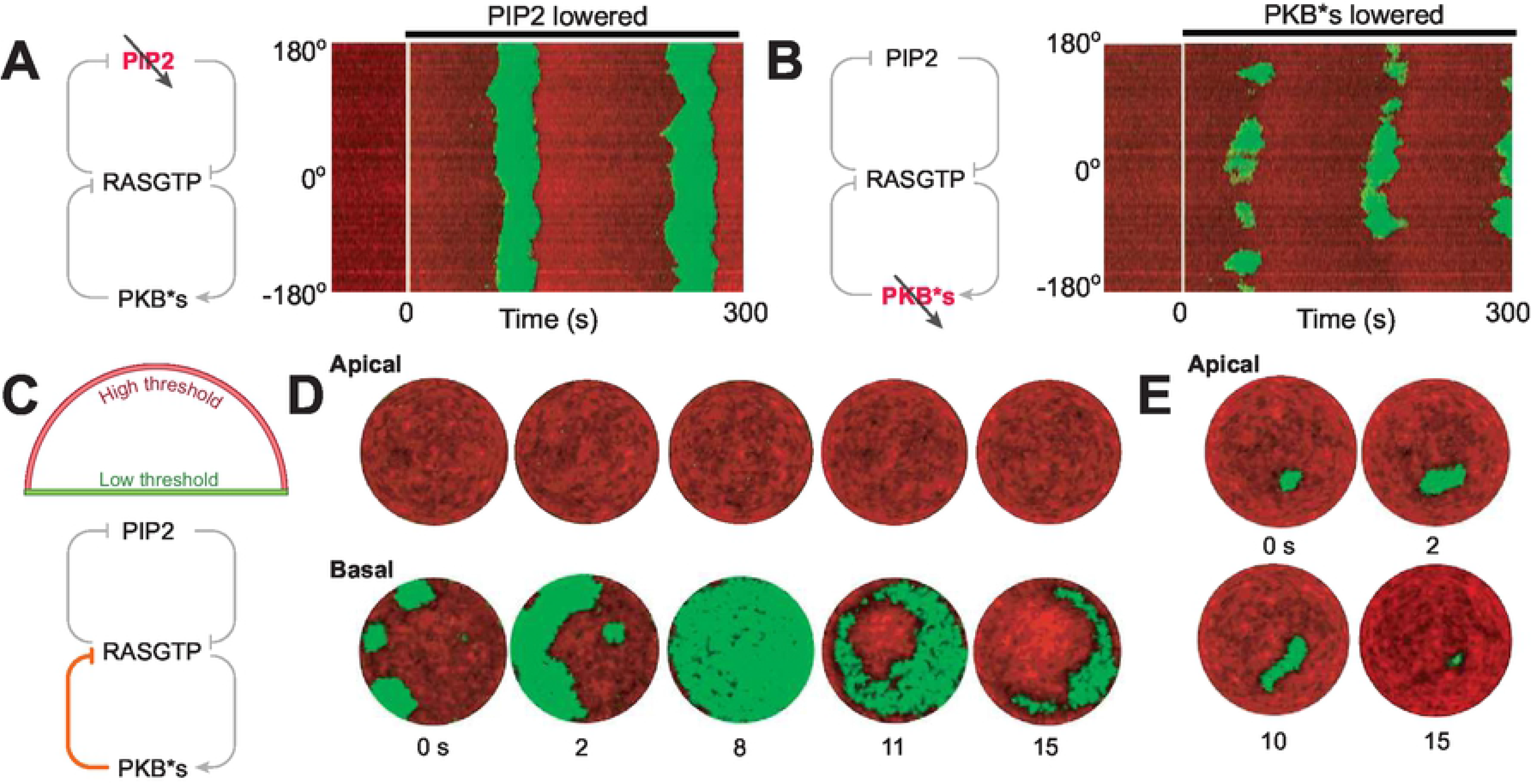
Effect of threshold on STEN dynamics. (A,B) The kymographs show the effect of lowering the STEN threshold by inhibiting PIP_2_(A) and PKB*s (B) as denoted in the schematics. The white lines indicate the time at which the respective species were lowered. (C) Schematic for incorporating the differential threshold between top and bottom surface of the cell through altering the PKB*s mediated inhibition on RasGTP. (D) Increased threshold restricts the wave activities at the basal surface and the waves were not allowed to travel to the apical surface. (E) Higher threshold on the apical surface made small waves with fewer number of wave initiations.

Experiments have also demonstrated that mechanical contacts can trigger excitable behavior [59–61]. Finally, we considered the possibility that the threshold was differentially regulated by mechanical contact with the substrate, with a lower threshold at the basal surface than at the apical surface. To take this into account, we assumed that the PKBs-mediated inhibition of RasGTP was different between the basal and apical surfaces, with lower inhibition at the former (Fig. 7C). In these simulations we observed frequent wave initiations at the basal surface, but these waves were quickly extinguished when they reached the apical surface (Fig. 7D). At the apical surface we rarely saw any de novo waves, and when they did appear, they did not spread like the basal counterpart and were extinguished rapidly (Fig. 7E).

## Discussion

Mathematical models have been a popular means of understanding how cells move in response to chemoattractant stimuli. These models have taken a modular view of the overall signaling network, breaking the overall system down into simpler functional blocks [62]. In doing so, most models have focused on a specific set of experimental observations, such as receptor-ligand interactions [35], adaptation [13, 16, 63], amplification [64], wave propagation [65–67], excitability [25, 47, 68], or polarity [69–71], by concentrating on a specific functional block sometimes ignoring its connection to the overall network. Part of this problem is a lack of experimental data as most experimental papers focus on specific aspects of the chemotactic behavior. Here, we have proposed an integrated model that takes the receptor-ligand binding to PKBs, which have been shown to be a link to the cytoskeletal network. Where the experiments do not offer single out a unique alternative, we have presented various possibilities and simulated scenarios that could be used to distinguish among them. Our model recreates most of the observed responses, including adaptation to persistent global stimuli (Fig. 5C–G), presence of secondary peaks (S7 FigC–F, [72]), spatially localized responses to gradients (Fig. 5H–I) along with the typical excitable behaviors such as wave annihilation and refractory period.

Because of the importance of noise-induced transitions in triggering the signal transduction excitable network, thereby allowing unstimulated cells to move randomly, our modeling approach has emphasized the role of noise. While the role of stochastic fluctuations in patterning cellular responses is now firmly appreciated, most of these studies consider molecules and species that exist in relatively small numbers [31, 32]. Our simulations here demonstrate, however, that even in signaling system that consist of biochemical species with high numbers, noise can play an important role in regulating cell physiology. The stochastic framework that we used to simulate our model does not require an artificial injection of noise. Instead, noise is a natural consequence of the stochastic description of the reaction-diffusion equations. In this stochastic setting, the LEGI mechanism adapts perfectly as in the deterministic setting, as long as we focus on the mean level of activity (Fig 3D, S4A,B). However, we showed that the variance in the response increases as a function of the stimulus strength (Fig. 3F) and that the size of this increase depends on the particular way that the LEGI is implemented. The increase in the variance observed could account for chemokinesis, the increased speed of random migration seen following a uniform chemokine stimulus in neutrophils [73]. Moreover, because the different implementations of the LEGI mechanism have distinctive noise patterns, our results suggest a way of elucidating the precise nature of the adaptation mechanism experimentally.

The biased-excitable network hypothesis suggests that chemoattractant stimulation leads to a signal that lowers the threshold of activation of the STEN, and various models and experiments support this view. However, how cells display a persistent crescent of front-markers towards the source of a chemoattractant gradient remains an open question [74]. Simulations of cells stimulated by gradients of chemoattractant show that the RR activity remains high at the cell front and is suppressed elsewhere. One would expect that the resultant spatially localized lowered threshold would trigger activity that would travel away from their point of origin. Further activity would be possible, after the delay imposed by the refractory period. To explain this, we modified the STEN dynamics such that, following stimulation, the system undergoes an excitable-to-bistable transition (S7 FigG,H). In this case, a persistent, elevated level of RR leads to a new “high” state of activity in the STEN, and persistent crescents were observed. Though our implementation is similar to the wave pinning scheme used to explain persistent polarization [75], it differs due to the fact that our system is only in the bistable region in the presence of a persistent high stimulus.

Our simulations show the power of having an excitable system at the heart of the chemotactic signaling system. This threshold becomes the most important parameter shaping the cellular response, dictating overall activity and eventually, migratory modes [43]. Increased number of wave initiations as well as more wave spreading is often associated with lowering of the threshold, whereas increase in the threshold corresponds to opposite effect. The threshold in the chemotactic signaling network can be altered in several ways and not all the alterations have the same effect on the system behavior. Theoretically weakening of the all the inhibitory pathways or strengthening the catalytic pathways acting on RasGTP can result lowering of threshold. Experimentally, several pharmacological/genetic perturbations have been used to study this effect of alteration in *Dictyostelium* cells [43, 44, 76]. Recent publications suggested of possible existence of a number of mechanical feedbacks on the leading edge protrusion [59, 77] and hence on the system threshold. Cao et al. reported that the basal surface waves are mostly restricted at the bottom surface unless the threshold of the system is lowered [60]. In that case, on reaching the bottom boundary the basal surface waves starts travelling upwards long the apical surface.

Finally, while our study has centered on chemoattractant signaling, we note that various components, such as GPCR signaling, the presence of feedforward adaptive mechanisms, and biochemical excitability are concepts that extend well beyond the realm of cell migration. For example, a recent report suggests that stochastic effects may play an important role in rhodopsin signaling [78]. Thus, our study provides a means for analyzing the effect of noise in these systems and could help to understand more complex interactions in these other settings.

## Supporting information

**S1 Fig. Effect of mesh refinement.** (A) Total number of nodes (in logarithmic scale) for different values of H_max_ (maximum allowable distance between nodes). H_min_ (minimum allowable distance between nodes) was chosen to be 1/3 of the respective H_max_ value. The red dot shows the mesh size adopted for the present study whereas the gray dots are used for the comparison in (C). (B) Plot of simulation time (in logarithmic scale) for different values of H_max_ corresponds to a one second simulation of a diffusion process involving a single entity. Colors are same as in (A). (C) Comparison of the simulation output (basal surface profile) of a diffusion process involving single entity at *t* = 1 s using fine, medium and coarse meshes as indicated by the H_max_ values. (D) Plot of simulation time (in logarithmic scale) for different values of H_max_ corresponding to a one second simulation of GPCR and LEGI modules combined, in the absence of cAMP. Colors are same as in (A).

**S2 Fig. Temporal response of different full- and reduced-order receptor states.** (A) Temporal responses of free (H,L,S), occupied (H:C,L:C,S:C), free phosphorylated (^P^H,^P^L) and occupied phosphorylated (^P^H:C,^P^L:C) states of the receptors, in absence and presence of a saturating dose of cAMP. The gray boxes denote the time segments used to compute steady-state concentration of different entities in (B). (B) Relative steady-state amount of different receptor states in absence (top) and presence (bottom) of cAMP. (C) Mean Fano factor of the temporal fluctuations among nodes for full-(blue solid) and reduced-order (red dashed) modules in response to varying cAMP levels (% R.O.). (D) Plot of total number of occupied receptors at nodes. Solid line and shaded region show the mean and the standard deviation respectively. (E) Comparison between internode noise characteristics of full-(solid) and reduced-order (dashed) modules in terms of the coefficient of variation (blue) and Fano factor (red). (F) Schematic of a 2-state reduced order model of receptors containing single unoccupied (R) and occupied (R:C) states. (G,H) Comparison between global responses of full-(blue solid) and reduced-order (red dashed) receptor modules in presence and absence of cAMP. The region where the two responses differ is denoted by the gray box in (G), which is zoomed into in (H).

**S3 Fig. G-Protein response and LEGI signaling.** (A) Dissociated G-protein response (%) to varying degree of receptor occupancy (% R.O.) with different total number of G-protein molecules (represented by different colors and denoted as a ratio to the total receptors molecules). (B) G-protein responses of (A) normalized to respective maximum values. The inset shows a zoomed-in version of a smaller section of the plot. Colors are same as in (A). (C) Coefficient of variation as a function of %R.O.. Colors are same as in (A). (D) Schematic of implicit LEGI mechanisms. (E,F) Comparison between implicit (left) and explicit (right) schemes of difference (C) and ratio (bottom) mechanism. Explicit mechanism involves the detailed reactions involving activator G_*βγ*_, inhibitor: I* and response regulator: RR, whereas the implicit schemes use the steady-state expression of RR directly in the model ignoring the transient dynamics. (G) Effect of concentrations of cAMP (in terms of % R.O.) on steady-state amplitude (blue), peak amplitude (red), adaptation time (green) and peak time (orange) of the nodal average profile of RR in the ratio mechanism for different basal activation/inactivation rate. The solid lines and the shaded regions show the respective mean and standard deviations (*n* = 10 independent simulations in which the parameter values were varied according to the distributions of Table 3).

**S4 Fig. Temporal response of LEGI mechanisms.** (A,B) Temporal nodal profiles of G_*βγ*_ (green), I^⋆^ (red) and RR (blue) in response to a staircase profile of cAMP for the difference (A) and ratio (B) mechanisms. (C,D) Temporal average (C) and individual (D) nodal profiles of G_*βγ*_ (green), I^⋆^ (red) and RR (blue) in response to the application and withdrawal of stimulus for difference (center) and ratio (bottom) mechanisms. The shaded regions in (C) denote standard deviations among all nodes from a single simulation.

**S5 Fig. Effect of diffusion of LEGI inhibitor during gradient stimulation.** (A,B,C) Temporal difference (between front and back of the cell) profiles of G_*βγ*_ (A), I^⋆^ (B) and response regulator (C, scaled) for different relative diffusive strengths of LEGI inhibitor, I^⋆^. The gray boxes denote the time segments used to generate the average values of the individual profiles for (D). (D) Plot showing steady-state temporal average values of the difference profiles G_*βγ*_ (green), I^⋆^ (red) and response regulator (blue) for different relative diffusive strengths of I^⋆^ (in logarithmic scale). Mean±SEM is shown (*n* = 5 independent simulations in which the parameter values are allowed to vary according to the distributions of Table 3).

**S6 Fig. Shallow gradient sensing by LEGI difference scheme.** (A) Schematic showing the diameter at the bottom surface (red) and a semicircular arc on the curved surface (blue) connecting front and back of the cell with respect to the needle position (light blue dot). (B) Circular cross sections of the hemispherical domain at *z* = 1 (a) and *z* = 2 μm (b). The front (closest to needle) and back of the cell is marked as 0 and 180°, respectively. (C) Temporal profile of G_*βγ*_ (green), I^⋆^ (red) and RR (blue) at cell front (0°, darker shade) and back (180°, lighter shade). The solid lines and the shaded regions show the respective mean and standard deviations (*n* = 10 independent simulations in which the parameter values are allowed to vary according to the distributions of Table 3) (D-G) Spatial response of the system for receptor occupancy (R.O.), G_*βγ*_, I^⋆^ and response regulator (RR). The kymographs (D) are based on the maximal projection of the hemispherical domain (nodes between a and b) of panel B. The white dashed line indicates the time instant when cAMP gradient was applied. Panel E shows the spatial profiles at the basal and apical surfaces at *t* = 3 min. Panel G shows the spatial profiles along the lines marked in panel A. Lines denote mean and the shaded regions standard deviations (*n* = 5 independent simulations in which the parameter values are allowed to vary according to the distributions of Table 3).

**S7 Fig. STEN dynamics.** (A) Spatiotemporal profiles of RasGTP-PKB*s (green-blue, top) and PIP2-PKB*s (red-blue, bottom) on the basal surface showing wave-traveling. The regions bounded by white dashed lines show the membrane regions with high RasGTP. (B) The Color-coded overlays show the progression of waves as a function of time and computation of the wave speed. (C) The nodal responses of RasGTP (green), PIP_2_ (red) and PKBs* (blue) at three random membrane nodes in presence of a global cAMP stimulus. (D) Basal surface profile of RasGTP (green) and PIP_2_ (red) in presence of a global stimulus at the time points indicated. (E,F) Normalized global response of RasGTP (D) and the corresponding kymograph in response to a sustained, spatially uniform cAMP dose. (G) Illustration showing how an excitable-to-bistable transition could occur. Shown are hypothetical nullclines for RasGTP (green) and PKB*s (blue) for a deterministic, two-state model for the excitable system. The darker and the lighter green shade represent the corresponding RasGTP nullcline in the excitable and bistable regimes, respectively. (H) Illustration showing the effect of a gradient stimulus. The dashed line represents the RasGTP-nullcline in the absence of a stimulus. The darker and the lighter green shade represent the RasGTP nullclines at the front and back of the cell, respectively, when subjected to cAMP gradient. The scale bars in (A,B,D) represents 2 μm.

**S1 File. Supplementary methods.**

**S2 File. MATLAB implementation.** Matlab scrips used for simulating the various models. The codes will be made available on acceptance.

**S1 Video. Wave traveling.** Movie showing various views of the wave traveling. Corresponds to Fig. 5C.

**S2 Video. Wave splitting.** Movie showing various views of the wave splitting. Corresponds to Fig. 5D.

**S3 Video. Wave annihilation.** Movie showing various views of the wave annihilation. Corresponds to Fig. 5E.

**S4 Video. Global stimulus.** Movie showing various views of the response to a global chemoattractant stimulus. Corresponds to Fig. 5F.

**S5 Video. Gradient stimulus.** Movie showing various views of the response to a gradient chemoattractant stimulus. Corresponds to Fig. 6A.

**S6 Video. Response to a successive application of gradient stimulii** Corresponds to Fig. 6B.

**S7 Video. Response to a gradient stimulation followed by a global stimulation** Corresponds to Fig. 6C.

### Acknowledgments

This work was supported, in part, by DARPA under contract number HR0011-16-C-0139 (P.A.I.). The authors would like to thank all members of the P.A. Iglesias (Johns Hopkins University), P.N. Devreotes and D. Robinson laboratories (Johns Hopkins University School of Medicine) for their valuable contributions in various points of the research process.

### Author Contribution

Conceptualization, D.B. with P.A.I.; Methodology, D.B. with P.A.I.; Software, D.B. with P.A.I.; Formal Analysis, D.B. Investigation, D.B. with P.N.D. and P.A.I.; Data Curation, D.B. Writing – Original Draft, D.B. with P.A.I.; Writing – Review & Editing, all authors; Visualization, D.B. with P.A.I.; Supervision, P.A.I.; Funding Acquisition, P.A.I.

